# Methyl viologen can affect mitochondrial function in Arabidopsis

**DOI:** 10.1101/436543

**Authors:** Fuqiang Cui, Mikael Brosché, Alexey Shapiguzov, Xin-Qiang He, Julia P. Vainonen, Johanna Leppälä, Andrea Trotta, Saijaliisa Kangasjärvi, Jarkko Salojärvi, Jaakko Kangasjärvi, Kirk Overmyer

## Abstract

Reactive oxygen species (ROS) are key signalling intermediates in plant metabolism, defence, and stress adaptation. The chloroplast and mitochondria are centres of metabolic control and ROS production, which coordinate stress responses in other cell compartments. The herbicide and experimental tool, methyl viologen (MV) induces ROS generation in the chloroplast under illumination, but is also toxic in non-photosynthetic organisms. We used MV to probe plant ROS signalling in compartments other than the chloroplast. Taking a genetic approach in *Arabidopsis thaliana*, we used natural variation, QTL mapping, and mutant studies with MV in the light, but also under dark conditions, when the chloroplast electron transport is inactive. These studies revealed a light-independent MV-induced ROS-signalling pathway, suggesting mitochondrial involvement. Mitochondrial Mn SUPEROXIDE DISMUTASE was required for ROS-tolerance and the effect of MV was enhanced by exogenous sugar, providing further evidence for the role of mitochondria. Mutant and hormone feeding assays revealed roles for stress hormones in organellar ROS-responses. The *radical-induced cell death1* mutant, which is tolerant to MV-induced ROS and exhibits altered mitochondrial signalling, was used to probe interactions between organelles. Our studies implicate mitochondria in the response to ROS induced by MV.

## Introduction

The study of reactive oxygen species (ROS) has transformed in the last decade, shifting our view from ROS as indiscriminate damaging agents to versatile and specific signal transduction intermediates. Plants have an enormous capacity to detoxify ROS, whose accumulation is rarely accidental, rather specific signalling events carefully orchestrated by the plant (Foyer & Noctor, 2016; Waszczak *et al.*, 2018). Due to ease of use, paraquat is a commonly used ROS generator for the study of ROS signalling. Paraquat is the common name of the herbicide methyl viologen (MV; N,-N′-dimethyl-4,-4′-bipyridinium dichloride), which acts in the production of ROS via a light dependent mechanism. In chloroplasts MV competes with ferredoxin for electrons on the acceptor side of Photosystem I (PSI; Dodge, 1989; Fuerst & Norman, 1991) and forms the MV cation radical, which reacts instantly with O_2_ to form superoxide (O_2_·^−^; Hassan, 1984). O_2_·^−^ subsequently forms other ROS and can cause cell death (Babbs *et al.*, 1989). This widely accepted view of MV as an inducer of toxic ROS is the relevant mechanism when used at high concentrations as an herbicide in the field. However, use at low concentrations as an experimental tool should be reconsidered in light of the current understanding of ROS signalling and processing.

Known MV tolerance mechanisms involve ROS detoxification, MV transport or sequestration, and chloroplast physiology (Vaughn *et al.*, 1989; Aono *et al.*, 1995; Van Camp *et al.*, 1996; Lasat *et al.*, 1997; Váradi *et al.*, 2000; Abarca *et al.*, 2001; Murgia *et al.*, 2004; Yu *et al.*, 2004; Davletova *et al.*, 2005; Miller *et al.*, 2007; Fujita *et al.*, 2012; Xi *et al.*, 2012; Li *et al.*, 2013; Hawkes, 2014). A relationship between long life span, sucrose availability, and tolerance against MV-induced ROS was seen in *gigantea* mutants (Kurepa *et al.*, 1998a) and exogenous sucrose treatment was shown to enhance MV toxicity (Kurepa *et al.*, 1998a, Kurepa *et al.*, 1998b), however the mechanism for this effect remains unknown. In *Arabidopsis* (*Arabidopsis thaliana*) forward genetic screens for MV tolerance mutants have yielded some insights into chloroplast ROS signalling (Chen *et al.*, 2009; Fujita *et al.*, 2012; Xi *et al.*, 2012; Fujita & Shinozaki, 2014; Luo *et al.*, 2016). RADICAL-INDUCED CELL DEATH1 (RCD1) was isolated as a ROS signalling component (Belles-Boix *et al.*, 2000; Overmyer *et al.*, 2000) and was found to alter tolerance to MV-induced ROS (Ahlfors *et al.*, 2004; Fujibe *et al.*, 2004). The RCD1 protein interacts with several transcription factors (Ahlfors *et al.*, 2004; Jaspers *et al.*, 2010) and functions as an integration point for multiple hormone and ROS signals (Jaspers *et al.*, 2009).

MV induces ROS production in all organisms tested, causing ROS production in mitochondria of non-photosynthetic organisms (Krall *et al.*, 1988; Minton *et al.*, 1990; Cochemé & Murphy, 2008). In plants, the induction of ROS signals by MV outside the chloroplast has been documented (Bowler *et al.*, 1991) but has remained mostly uncharacterized. Many studies have used MV treatment to test general ROS responses; however, few of these directly used MV as a tool to address ROS or redox signalling and their associated pathways. Thus, we used MV as a tool under both light and dark conditions to probe the genetics of ROS responses in and outside the chloroplast. We show an important function for mitochondria in ROS signalling induced by low concentration MV-treatment in the dark.

## Materials and Methods

### Plant material and growth

*Arabidopsis* (*Arabidopsis thaliana*) genetic resources were obtained from NASC (www.arabidopsis.info). All mutants were PCR genotyped and confirmed over two generations. Double mutant construction has been presented elsewhere (Brosché *et al.*, 2014), primers used in genotyping mutants are listed in Table S1.

Aseptic cultures were performed on 135 mm square plates in the presence or absence of MV as indicated, on 0.5x MS (Murashige and Skoog) medium containing 0.8% agar, 0.05% MES (pH 5.7) and 1% sucrose, except as otherwise noted. Following a three-day stratification (4°C in the dark) seeds were light treated for 4 hr to promote germination and then placed vertically in an environmental chamber (Sanyo; www.sanyo-biomedical.co.uk) with 12/12 hr day/night cycle, constant 20°C, and light of 120 μmol of photons m^−2^ s^−1^. For dark treatments, plates were covered with two layers of aluminium foil.

### Growth and chlorophyll fluorescence assays

For growth measurements, eight- or nine-day-old seedlings were photographed with a size scale then hypocotyl- or root-lengths were determined with ImageJ software (http://rsbweb.nih.gov/ij/). Chlorophyll fluorescence imaging was performed as described (Barbagallo *et al.*, 2003); briefly, 1-2 seeds were sown in each well (with 0.180 ml media) of a black 96-well-plate and sealed with plastic film. Seedlings were grown under standard conditions with 220 μmol of photons m^−2^ s^−1^ for four or five days before treatments. MV was added to a final concentration of 250 μM. All plates were placed in the dark for 20 minutes and then were placed in the light (160 μmol of photons m^−2^ s^−1^) for 6-8 hr or in the dark for 20 hr before measurements. Salicylic acid, methyl jasmonate, abscisic acid, and 1-aminocyclopropane-1-carboxylic acid (ACC) (Sigma; www.sigmaaldrich.com) were added to a final concentration of 200 μM 14 hr prior to MV for hormone protection experiments. Whole plate imaging utilized a Walz M-series imaging PAM Chlorophyll fluorescence system (www.walz.com) using the maxi head. Measurement of quantum efficiency of PSII (F_v_ F_m_^−1^) from individual wells was then calculated with Walz Imaging Win software. Before measurements, seedlings were dark adapted for 20 minutes.

### H_2_ O_2_ staining

H_2_ O_2_ accumulation was visualized by staining with 1mg/ml 3,3’-diaminobenzidine (DAB) in 10 mM NaHPO_4_ (pH 4.0). Detached rosettes of 18-day-old soil grown Col-0 and *rcd1* plants were floated on water (ddH_2_O with 0.05% Tween20), or water containing 1μM MV, overnight (15 hrs) in the dark. Plants were then pre-treated for 0-2 hrs in the light (250 μmoles m^−2^ sec^−1^), before vacuum infiltration with DAB and stained for 5 hrs in the dark. Samples were fixed and cleared in 95% ETOH: 85% lactate: glycerol (3:1:1) for 2-10 days. Cleared samples were stored and mounted in 60% glycerol.

### Light treatments

For photoinhibition under high light, 11-day-old plate-grown seedlings were placed in the imaging PAM chlorophyll fluorescence system and subjected to intermittent high light, consisting of 60-minute illumination with strong blue light (200 μmol of photons m^−2^ s^−1^), 25 minutes of darkness, then F_0_ and F_m_ were registered, after which the next cycle began. To avoid overheating, continuous cooling to room temperature was used by running tap water through coiled rubber tubing beneath. Photoinhibition was observed as decreased F_v_ F_m_^−1^ = (F_m_ – F_0_) F_m_^−1^. For fluctuating light treatments, plants were grown on soil with an alternating 5 min low light (50 μmol photons m^−2^ s^−1^) and 1 min high light (500 μmol photons m^−2^ s^−1^) illumination (Tikkanen *et al.*, 2010) throughout the entire 8 hr light period of an 8/16 h light/dark cycle).

### Chlorophyll measurements

Leaf disks (7 mm) from the first two fully expanded middle-aged leaves were infiltrated with 0.5x MS liquid with MV and placed on similar MV containing solid media plates for 14 hr under light or dark condition before photographing. Pigments were extracted in 80% acetone and absorbance measured at 645 and 663 nm using a spectrophotometer (Agilent 8453; www.home.agilent.com). The total chlorophyll concentration was calculated using Arnon’s equation (Arnon, 1949).

### QTL mapping

The mapping population of 125 Kondara x L*er* recombinant inbred lines (RILs) was treated with or without 0.1 μM MV for growth assays or 250 μM MV for the fluorescence assay. For mapping the QTL in light/dark, the ratio of each line was obtained by using the mean of the root (in light) or hypocotyl (in dark) lengths of treated plants divided by control. For fluorescence assays, the F_v_ F_m_^−1^ of the controls were all the same, thus F_v_ F_m_^−1^ values after MV treatment were used directly for QTL mapping. Data normality was checked with quantile-quantile plots in R (R Development Core Team, 2014). Data for dark-grown seedlings was normally distributed but light grown was log_10_ transformed to gain normality. QTL mapping was performed with single-locus QTL scans with interval mapping. Chlorophyll fluorescence data could not be transformed to gain normality and therefore nonparametric interval mapping was conducted. The genome-wide LOD threshold for a QTL significance (P < 0.05) was calculated separately for each trait by 10,000 permutations. All the QTL analyses used R with R/qtl (Broman *et al.*, 2003).

### qPCR

Five-day-old *in vitro* grown seedlings were transferred to medium with or without 0.1 μM MV and collected two days later in liquid nitrogen for RNA extraction. Four-week-old soil grown plants were collected for RNA extraction (GeneJET Plant RNA Purification Mini Kit, Thermo Scientific). Reverse transcription was performed with 3 μg DNAseI treated RNA using RevertAid Premium Reverse Transcriptase (Thermo Scientific). The cDNA was diluted to 100 μl final volume. Three technical repeats with 1 μl cDNA and 5x HOT FIREPol EvaGreen qPCR Mix (Solis Biodyne) were used for qRT-PCR. Primer sequences and amplification efficiencies determined with the Bio-Rad CFX Manager program from a cDNA dilution series are given in Table S1. The raw cycle threshold values were analysed in Qbase+ (https://www.qbaseplus.com/; Hellemans *et al.* 2007) using *YLS8* (AT5G08290), *TIP41* (AT4G34270) and *PP2AA3* (AT1G13320) as the reference genes as described (Brosché *et al.*, 2014).

### Statistics

The statistical significance of the relative change in hypocotyl and root lengths was estimated using scripts in R. First, a logarithm of the raw hypocotyls length data was taken and a linear model was fitted with genotype, treatment, and their interaction terms. Model contrasts and their significances were estimated with multcomp package in R (Version 3.03; Bretz *et al.*, 2010). All experiments were repeated at least three times.

### Protein extraction and immunoblotting

Total proteins were extracted by grinding of frozen seedlings in RIPA buffer (50 mM Tris-HCl, pH 8.0, 150 mM NaCl, 1% Triton X-100, 0.5% sodium deoxycholate, 0.1% SDS) in the presence of protease inhibitor cocktail (Sigma-Aldrich; www.sigmaaldrich.com). The samples were centrifuged at 16,000 x g for 15 min and the supernatant used for western blotting. Protein concentration in the extracts was determined by Lowry method using the DC protein assay (BioRad; http://www.bio-rad.com).

Proteins (5 to 10 μg per lane) were separated using 15% SDS-PAGE gels in presence of 6 M urea and transferred onto PVDF membranes (BioRad). The membranes were blocked in 3% BSA in TBS-T (20 mM Tris-HCl, pH 7.5, 150 mM NaCl, 0.05% Tween-20) buffer and probed with an ASCOBATE PEROXIDASE (APX)-specific antibody diluted 1:2000 with TBS-T buffer containing 1% BSA. Horseradish peroxidase-conjugated donkey anti-rabbit IgG (GE Healthcare; www.gehealthcare.fi) was used as a secondary antibody and the signal was visualized by SuperSignal West Pico luminescence reagents (ThermoFisher Scientific; www.fishersci.fi).

### Abundance of photosynthetic complexes by 1-dimentional acrylamide gels

Fourteen-day-old seedlings from plates +/− MV (0.4 μM) were snap-frozen in liquid nitrogen and ground with glass beads in Precellys 24 tissue homogenizer (3 × 10 seconds at 6800 rpm). Total protein was extracted by incubation of the homogenate in 100 mM Tris (pH 7.8), 2% SDS, 1 × Protease Inhibitor Cocktail for 30 minutes at 37 °C. Protein samples were loaded on equal chlorophyll basis (0.45 μg of chlorophyll per well) and separated in 12 % acrylamide gels. Immunoblotting was performed with the antibodies raised against PSI subunit PsaB, PSII subunit PsbD, or LhcA2 and LhcB2 antennae proteins (Agrisera; www.agrisera.com).

## Results

### The dark response to MV

This study utilizes the MV tolerant *rcd1* mutant and its moderately tolerant Col-0 parental accession. Decreased expression or activity of MV transporters excludes MV from its active sites leading to stress avoidance (reviewed in Fujita & Shinozaki, 2014). To address this in the *rcd1* mutant, the expression of known MV transporters was tested. Only minor differences in expression between *rcd1* and Col-0 were observed and accumulation of the major plasma membrane importer, *PDR11*, was higher in *rcd1* (Fig. S1). These data suggest that *rcd1* did not avoid stress due to altered MV transport. Further, the effect of MV on PSI oxidation and initial H_2_O_2_ production was similar in Col-0 and *rcd1* (Shapiguzov *et al.*, 2018). Together this indicates that *rcd1* tolerance is not based on restricted access of MV to PSI. Thus, we use the *rcd1* mutant here as a tool to dissect MV-induced ROS signalling. Plant MV responses are dependent on light, growth, and assay conditions, which prompted us to evaluate these parameters. The response to MV-induced ROS was assayed *in vitro* on MS plates under standard light conditions (100 μmoles m^−2^ s^−1^) scored by visual appearance (Fig. 1). Root length was quantified in light-grown seedlings (Fig. 1b). Growth inhibition assays of four independent *rcd1* alleles (Jaspers *et al.*, 2009) indicated all were equally tolerant (Fig. S2a). The *rcd1*-1 allele was used in further experiments, hereafter referred to as *rcd1*. Three-week-old soil grown plants were assayed for leaf disk chlorophyll bleaching (Fig. 2) and in seven-day-old *in vitro* grown seedlings decreases in quantum efficiency of photosystem II (PSII) (F_v_ F_m_^−1^) was monitored as a stress index (Barbagallo *et al.*, 2003) using chlorophyll fluorescence (Fig. 2c). All assays detected differential tolerance to MV-induced ROS over a wide but variable range of concentrations. Root length (Fig. 1a,b) was the most sensitive assay detecting differences in the low nM range. The root length assay exhibited light intensity dependent effects of MV (not shown) and has been previously shown to correlate well with other light based assays, such as photosynthesis rate, leaf growth, and leaf chlorophyll bleaching (Davletova *et al.*, 2005; De Clercq *et al.*, 2013), thus was used here in subsequent studies of MV-induced ROS responses in the light.

**Fig. 1.**
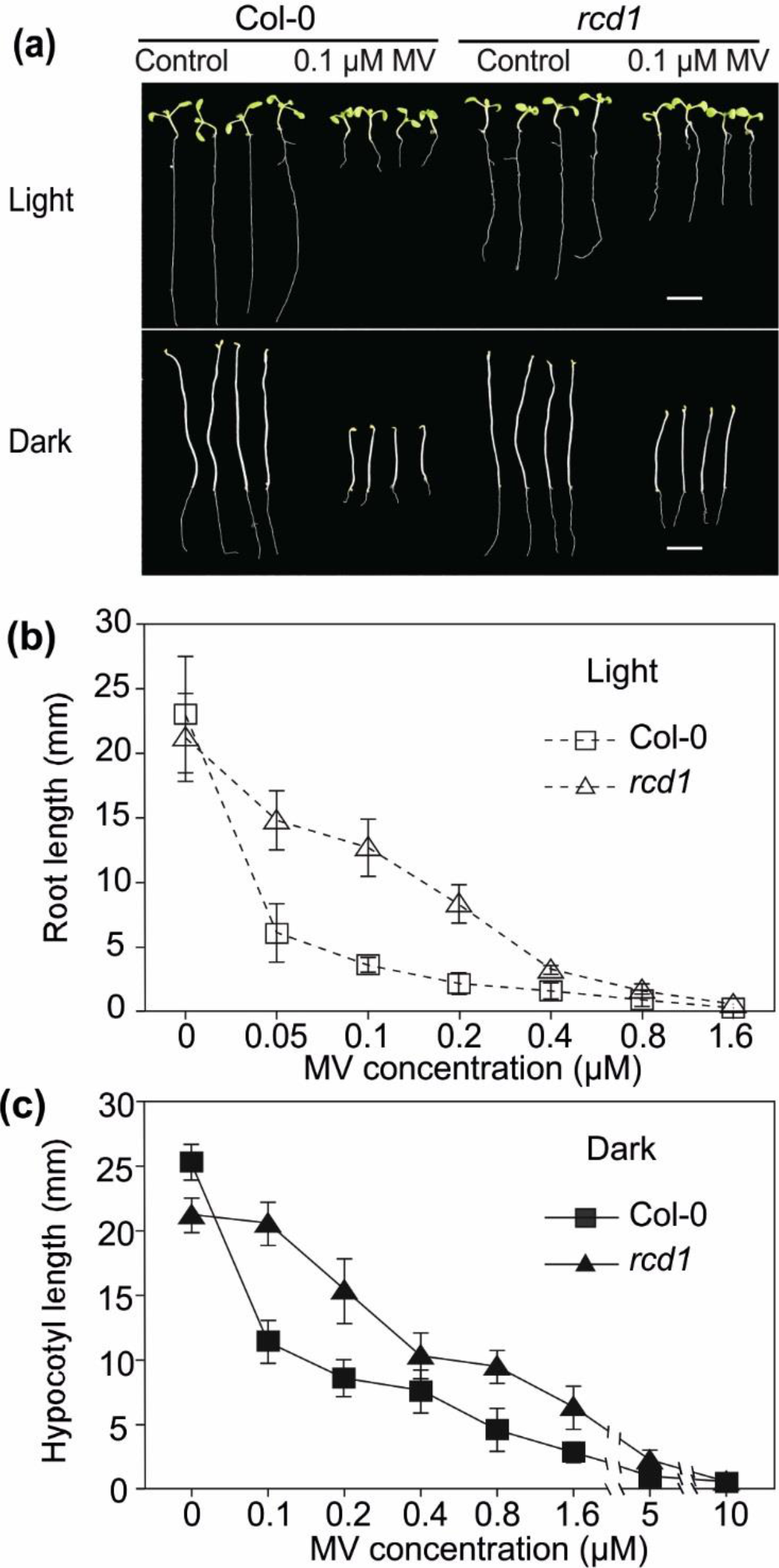
Methyl viologen (MV)-induced growth inhibition in light and dark. (a) Wild type (Col-0) and the *radical-induced cell death1* (*rcd1*) mutant after eight or nine days of growth on 0.1μM MV or control plates. Scale bar = 1 cm, N=12. (b) Quantification of root length in the light (c) or hypocotyl length in the dark at different MV concentrations. Results are presented as means ±SD (N=15). Wild type Col-0 and *rcd1* were grown eight or nine days in the light or dark on plates containing MV at the indicated concentrations, they were photographed and root or hypocotyl lengths quantified using ImageJ. All experiments were repeated three times with similar results and one representative experiment is shown

**Fig. 2.**
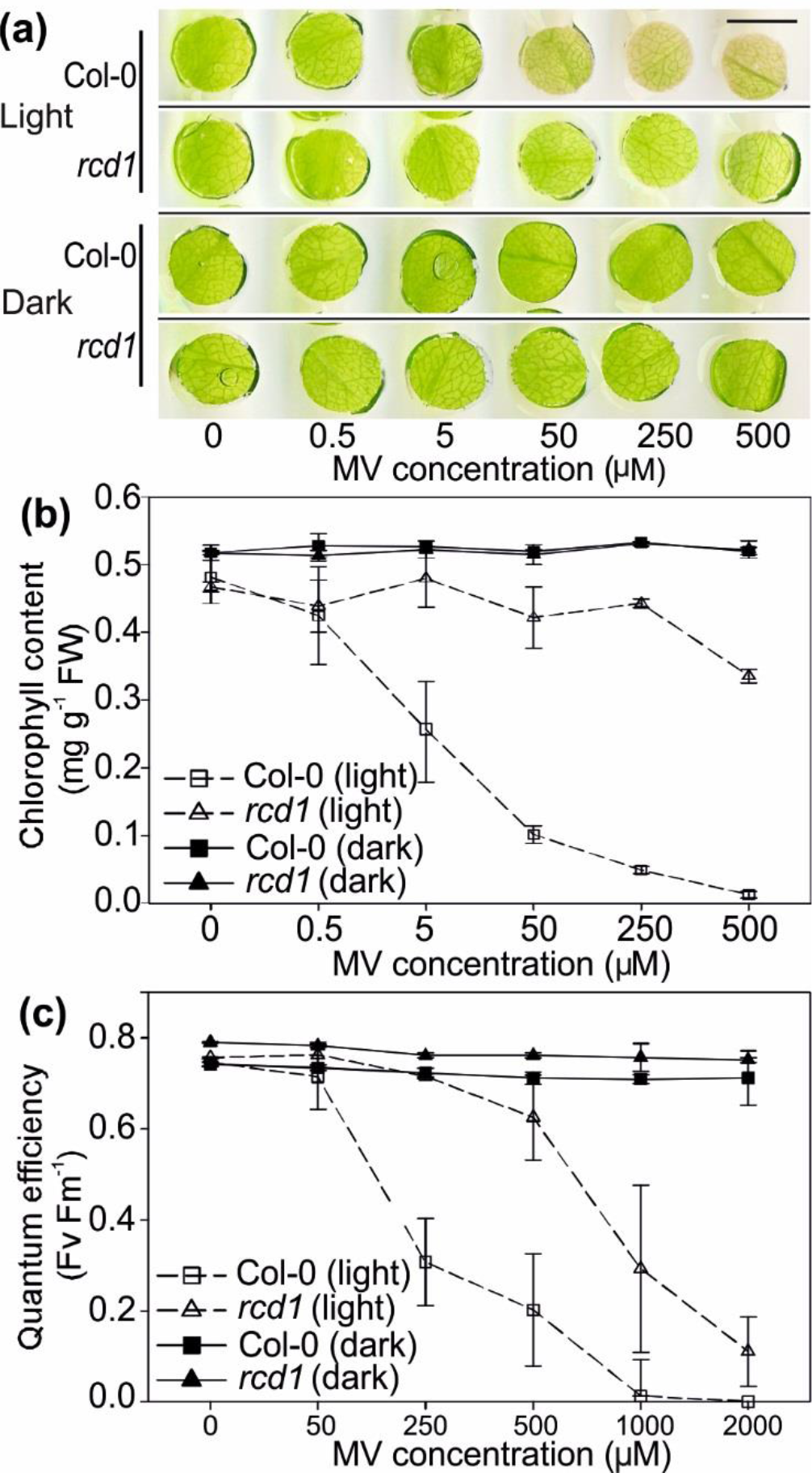
Methyl viologen (MV)-induced chloroplast damage in light and dark. (a) Leaf disks cut from three-week-old soil grown wild type (Col-0) and *radical-induced cell death1* (*rcd1*) mutant plants were treated *in vitro* with different concentrations of MV in light or dark for 16 hr, showing chlorophyll loss. Experiment was preformed three times with the same results, one representative experiment shown. Scale bar = 7 mm. (b) Quantification of chlorophyll content at different MV concentrations. Results are presented as means ±SD (N=4). Experiment was repeated three times with similar results. One representative experiment is shown. Chlorophyll content was determined spectrophotometrically from pigments extracts of leaf disks from plants grown and treated as in panel (a). (c) Quantum efficiency of photosystem II (F_v_ F_m_^−1^) measured by chlorophyll fluorescence at different MV concentrations. Results are presented as means ±SD (N=8). Experiment was repeated three times with similar results. One representative experiment is shown. Chlorophyll fluorescence was measured with an Imaging PAM from one-week-old seedlings, two per well in a 96 well plate containing 180 μl of 0.5x MS media treated with 20 μl stock solutions to give the indicated final concentrations of MV

To explore a potential role for non-photosynthetic processes in MV-induced ROS signalling, we assessed MV-induced ROS sensitivity in darkness, when photosynthetic electron transfer is inactive. Hypocotyl length was used as an index of MV-induced growth inhibition under dark conditions. MV inhibited hypocotyl elongation in both Col-0 and *rcd1* seedlings in the dark and the tolerance of *rcd1* was also observed here (Figs. 1a, c, S2b). In the dark, MV-induced changes were only detectable in growth-based assays. Chloroplast damage based assays exhibited no change by MV treatment in dark conditions (Fig. 2a-c).

To detect potential ROS sourced outside the chloroplast, we monitored MV-induced H_2_ O_2_ accumulation by DAB staining in the dark. Detached whole rosettes were loaded with 1 μM MV overnight in darkness, exposed to a two-hour light pulse, then transferred back to darkness for infiltration and staining with DAB for 5 hrs. In this experimental design, DAB is never present in the light. Col-0 plants exhibited marked accumulation of DAB precipitate (Fig. 3); importantly, this revealed accumulation of H_2_O_2_ in the darkness, when the chloroplast electron transfer chain is inactive. MV-tolerant *rcd1* mutant plants exhibited little change over the background stain intensity. This response in Col-0 plants was triggered by the light pre-treatment (Fig. S3). This indicated that MV-induced responses were initiated in chloroplasts, but the subsequent ROS production did not require active chloroplast electron transport.

**Fig. 3.**
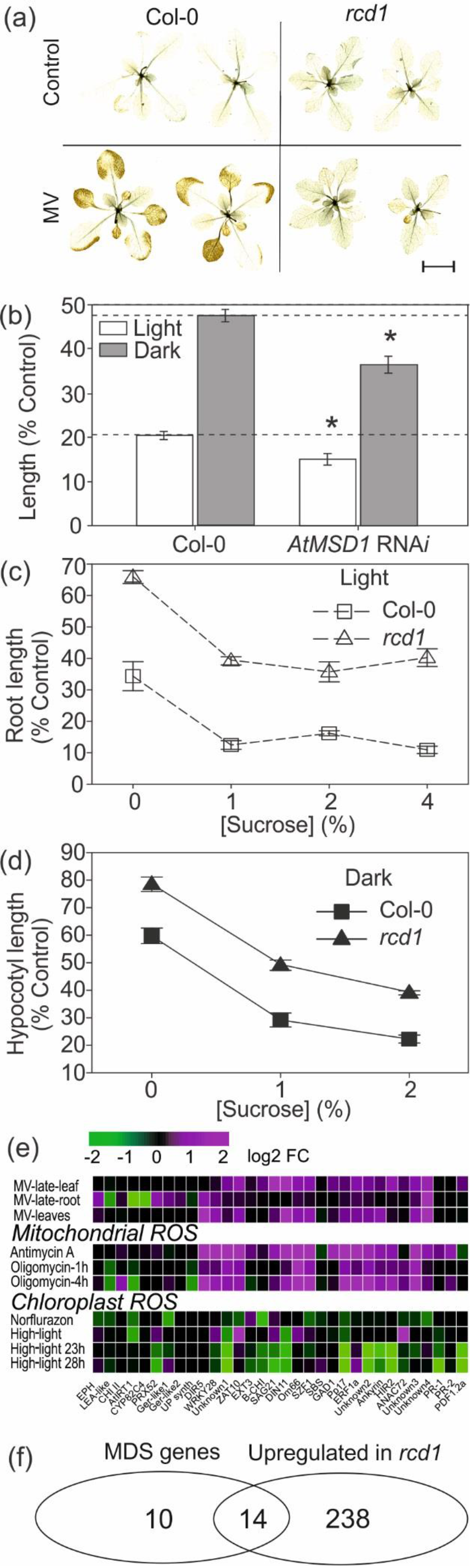
The involvement of mitochondria in methyl viologen (MV) toxicity. (a) H_2_ O_2_ accumulation in wild type (Col-0) and the *radical-induced cell death1* (*rcd1*) mutant induced by 1 μM methyl viologen (MV) and a 2 hr light pre-treatment as visualized with 5 hrs of diaminobenzidine (DAB) staining in the dark. Experiments repeated two times with similar results and one representative experiment shown. Separate images of stained plants are composited and separated by black lines. Images cut from one single photo of all treatments, which is presented in Fig. S4. Black scale bar = 1 cm and is valid for all images. (b) MV-induced ROS sensitivity of mitochondrial *MnSOD* silenced RNA*i* lines. Root length in the light or hypocotyl length in the dark were quantified from eight or nine days old seedlings grown on 0.5x MS plates (with 1% sucrose) with or without 0.1 μM MV and the results are presented as means ±SD (N=30) of the ratio between MV treated to control expressed as a percentage (% control). Experiment was repeated three times and results were pooled and analysed. Asterisks show statistical significance (P<0.05) from *post-hoc* analysis by computing contrasts from linear models and subjecting the P values to single-step error correction. (c) Quantification of root length in the light (d) and hypocotyl length in the dark at 0.1μM MV with different sucrose concentrations. Results are presented as means ±SD (N=24) of the ratio between MV treated to its respective control with the same sucrose concentration, expressed as a percentage (% control). Experiment was repeated three times and results were pooled and analysed using posthoc analysis by computing contrasts from linear models and subjecting the P values to single-step error correction. Measurements were taken from eight- or nine-day-old seedlings grown on 0.5x MS plates containing the indicated concentration of sucrose and 0.1μM MV. (e) Heat map depicting the expression of MV-response genes in the following treatments: methyl viologen (MV), inhibitors of mitochondrial function (Antimycin A and Oligomycin) and chloroplast stress (Norflurazon and high light). For comparison, the marker genes *PR-1*, *PR-2* (SA) and *PDF1.2* (JA) were also included. Magenta indicates increased expression and green decreased expression. The full gene list with AGI codes can be found in Table S2a. (f) Overlap between RCD1-regulated genes and mitochondrial dysfunction stimulon (MDS) genes regulated by ANAC013. Genes regulated downstream of RCD1 are from Jaspers *et al* (2009). MDS genes are as defined by De Clercq *et al.* (2013). List of genes, AGI codes, and their functional descriptions can be found in Table S2b

This was further addressed using genotypes or conditions known to enhance mitochondrial ROS accumulation. First, *AtMSD1* RNA*i* plants lacking the mitochondrial MnSOD, and thus deregulated mitochondrial ROS accumulation (Morgan *et al.*, 2008), were assayed. Under both light and dark conditions, *AtMSD1* RNA*i* plants exhibited enhanced growth inhibition by MV-induced ROS (Fig. 3b). Second, exogenous sugar increases oxidative phosphorylation and mitochondrial electron transfer (Fernie *et al.*, 2004; Keunen *et al.*, 2013), which could enhance ROS production by MV. Accordingly, such treatment was shown to enhance MV responses (Kurepa *et al.*, 1998b). To test this under conditions that control for any possible osmotic or sugar signalling effects, we used an experimental design that compensated for these effects by expressing the results as a ratio where plants treated with MV and sugar are normalized to respective control plates containing the same sugar concentration, but no MV. Exogenous sugar enhanced the inhibition of growth by MV both in the light and dark (Fig. 3c,d) suggesting that mitochondria are involved in MV action also under light. This effect was similar for sucrose (Fig. 3c,d) and glucose (Figs. S4, S5). Taking these results into account, additional MV dose response curves under different sugar concentrations (Figs. 3, S5), were used for selecting experimental conditions; unless otherwise indicated, 0.1 - 0.2 μM MV and 1% sucrose were used for all further experiments presented below.

### MV-induced mitochondrial signals

Additional support for the involvement of signals originating from mitochondria in MV responses was obtained from gene expression meta-analysis with data from Genevestigator (Hruz *et al.*, 2008). The expression of MV responsive genes was plotted in response to MV, inhibitors of mitochondrial function, and light treatments. This gene set was stringently defined and was previously found to be expressed in both photosynthetic and non-photosynthetic tissues, i.e. leaves and roots, treated with MV (Hahn *et al.*, 2013). Transcript abundance of these genes was higher in response to both MV and mitochondrial inhibitors, but lower in response to high light (Fig. 3e, Table S2a).

Analysis of genes deregulated in the MV-tolerant *rcd1* mutant provides further evidence of mitochondrial involvement. RCD1 is known to interact with transcription factors that control expression of mitochondrial dysfunction stimulon (MDS) genes (Jaspers *et al.*, 2009; Van Aken *et al.*, 2009; De Clercq *et al.*, 2013; Shapiguzov *et al.*, 2018). MDS genes are nuclear encoded genes for mitochondria localized proteins that are transcriptionally activated via mitochondrial retrograde regulation (MRR) upon the disturbance of mitochondrial function by stress. A clear overlap and statistically significant enrichment is seen when genes deregulated in *rcd1* are compared with MDS genes (Fig. 3f, Table S2b; Cluster IIIb in Brosché *et al.*, 2014). Together, these findings support that RCD1 regulates mitochondrial processes.

### Chloroplast-mitochondrial interactions in MV response

Loss of RCD1 function results in marked alterations in mitochondrial functions (Shapiguzov *et al.*, 2018). However, the question remains unresolved to which extent mitochondria contribute to chloroplast-related phenotypes of *rcd1* including tolerance to MV-induced ROS. To address this, we quantitatively tested the *rcd1* mutant for tolerance to chloroplast stress induced by high light (Fig. 4). Plant stress levels were monitored by measuring F_v_ F_m_^−1^ between pulses of high light (1200 μmol of photons m^−2^ s^−1^) over a 12 hr time course. The *rcd1* mutant reproducibly exhibited only slightly lower PSII photoinhibition levels throughout the entire 12 hr experiment (Fig. 4a), thus *rcd1* exhibits only a low level of tolerance to high light. To further test this we utilized the genes that are deregulated in *rcd1*, which we previously identified (Jaspers *et al.*, 2009) and queried against databases of experimentally determined chloroplast and mitochondria resident proteins using fisher’s exact test to discern enrichment for proteins localized to these organelles. Target genes downstream of RCD1 exhibited a significant enrichment (p=0.0008544) for genes encoding mitochondria localized proteins, but no enrichment (p=0.08316) for genes encoding chloroplast proteins. These results further support that RCD1 regulates primarily mitochondrial processes. Thus, we concluded that the reasons for physiological abnormalities observed in *rcd1* are of predominantly mitochondrial origin.

**Fig. 4.**
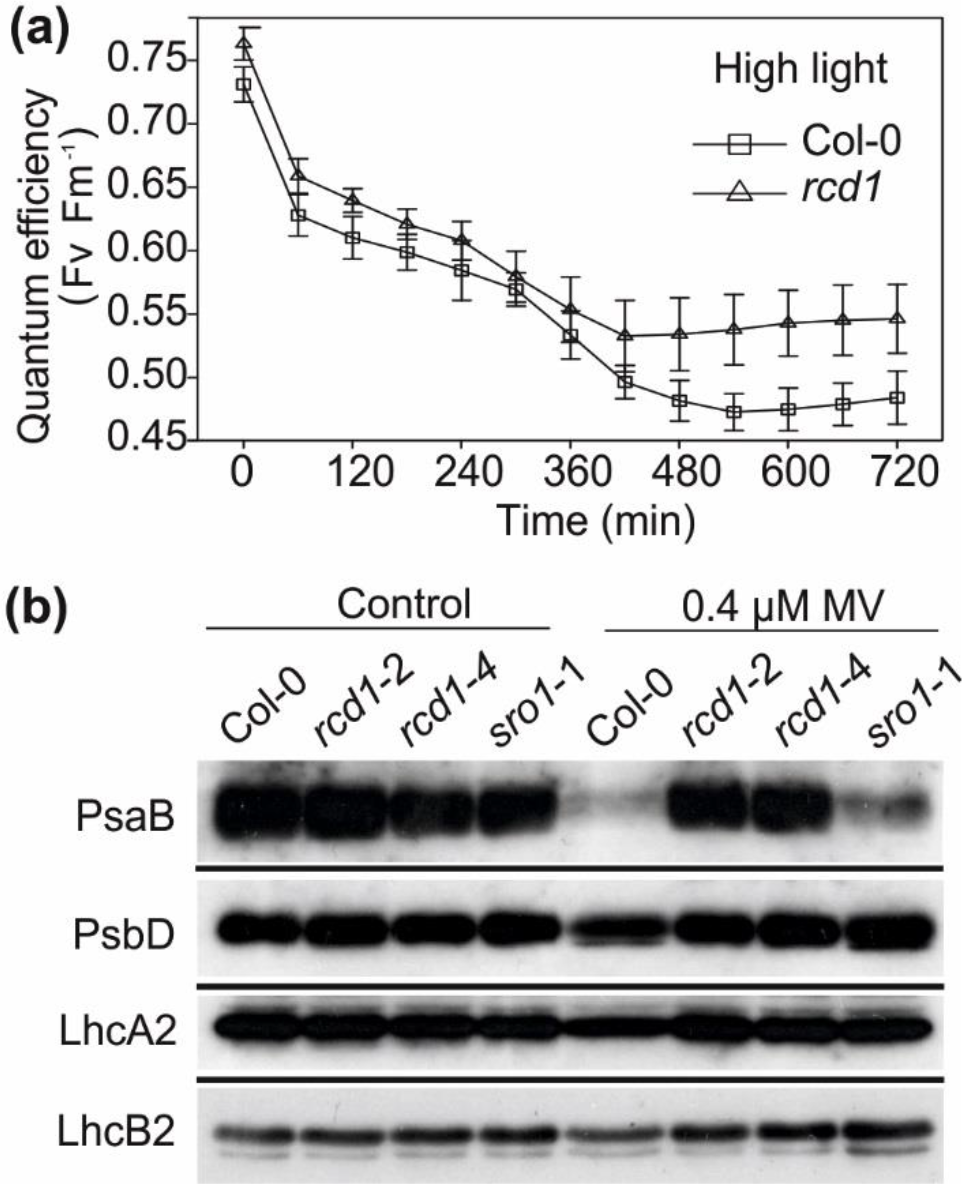
Chloroplast adaptation to ROS-inducing treatments. (a) Time course of high light induced decreases in quantum efficiency (F_v_ F_m_^−1^) of photosystem II (PSII); measured by chlorophyll fluorescence in plants exposed to repeated pulses of high light (1200 μM photons m^−2^ s^−1^ for 60 min followed by a 25 min dark adaptation) in wild type (Col-0) and *radical-induced cell death1* (*rcd1*) mutant plants. Chlorophyll fluorescence was measured with an Imaging PAM from one-week-old seedlings. Results are expressed as means ± SD (n=15), four biological repeats were done and one representative experiment is shown. (b) Abundance of chloroplast photosynthetic complexes as determined by protein immunoblotting. Seedlings were germinated and grown on MS plates with or without 0.4 μM methyl viologen (MV). Total protein extracts loaded on equal chlorophyll basis were separated by SDS-PAGE and blotted with anti-PsaB and anti-PsbD antibodies to assess the amounts of PSI and PSII, accordingly. Light harvesting antennae were analysed with anti-LhcA2 and anti-LhcB2 antibodies. The RCD1 paralog SIMILAR TO RCD-One (SRO1) is included here as a control. Experiment was performed twice with the same results with one representative experiment shown

Given the known coordination between the mitochondria and chloroplasts in metabolism and energy production (Noguchi & Yoshida, 2008; Vanlerberghe *et al.*, 2016), we next used the *rcd1* mutant to probe the interaction of mitochondrial and chloroplastic ROS processing systems. For this, the abundance and configuration of photosynthetic machinery was tested in Col-0 and *rcd1* under severe light stress conditions. Plants were grown under fluctuating light (constant alternation between 5 min low light and 1 min high light illumination during the entire day period; Tikkanen *et al.*, 2010). Thylakoid membrane protein complexes were isolated and separated on 2D gels utilizing a blue native gel in the first dimension and SDS PAGE in the second. This revealed increased abundance of PSI supercomplexes in *rcd1* under fluctuating light (Fig. S6a,b), suggesting the effect of RCD1 and possibly mitochondria on regulation of PSII to PSI stoichiometry in the chloroplasts. In particular, maintenance of PSI was affected. PSI is the primary target of MV under light, thus regulation of its abundance was tested under MV stress conditions. Col-0 seedlings germinated and grown in the presence of MV contained less chlorophyll than *rcd1* (Fig. 2a,b). To compensate for this, protein extracts from MV-treated Col-0 and *rcd1* were loaded on the gel on equal chlorophyll basis (Fig. 4b), Col-0 seedlings displayed dramatically decreased PSI levels (judged by abundance of the core protein, PsaB) vs. PSII (PsbD) or light-harvesting antenna (LhcA2 and LhcB2). This MV-dependent decrease in PSI was absent from the *rcd1* mutant (Fig. 4b). Thus, the stoichiometry of photosynthetic complexes was affected by development in the presence of MV in the wild type, but not in *rcd1*. Together, these findings suggested that adjustments of the photosynthetic apparatus under light stress was dependent on RCD1 function.

### Cytosolic APX in MV-triggered ROS responses

Fujibe *et al.* (2004) reported higher chloroplast stromal APX (sAPX) and chloroplast thylakoid APX (tAPX) transcript accumulation in the *rcd1* mutant, suggesting its tolerance to MV-induced ROS was due to enhanced ROS detoxification. Further, it has been proposed that APXs have a significant role in regulating tolerance to MV-induced chloroplast ROS (Davletova *et al.*, 2005). We utilized mutants with APX function compromised in specific compartments; the cytosolic cAPX1 and two in the chloroplast, sAPX and tAPX. Mutants were confirmed by protein immunoblot to be protein null (Fig. S7a,b), including a new allele of the *capx1* mutant in the Col-0 genetic background (SAIL_1253_G09), here designated as *capx1-2*. The *capx1-2* mutant exhibited enhanced growth inhibition by MV-induced ROS both in the light and dark, while the *sapx, tapx* single- and *sapx tapx* double mutants behaved as wild type under all conditions (Fig. S7c). The reduced growth observed in soil-grown *capx1-1* (Ws-0) under normal growth conditions (Davletova *et al.*, 2005), was not observed in *capx1-2* in Col-0 background (Fig. S7d). No differences in the protein levels of cAPX, tAPX or sAPX were observed in the *rcd1* mutant (Fig. S7a). These results implicate cAPX, but suggested that MV-induced ROS tolerance of RCD1 can not be explained by the accumulation of chloroplast-localized APXs, prompting further genetic experiments to explore other mechanisms.

### Natural variation of MV response

Our data implicating mitochondria in the MV-induced ROS response relies entirely on a single accession of *Arabidopsis* (Col-0). To seek additional evidence, natural variation in the organellar ROS sensitivity of 93 diverse accessions (Nordborg *et al.*, 2005) of *Arabidopsis* was surveyed. This was first performed in the light using three different assays. A plate germination screen with 0.5 and 1.0 μM MV was visually scored based on growth using a scale of 1-4 (Fig. S8a). Root growth and PSII quantum efficiency were used as quantitative assays (Fig. S8b,c). The *rcd1* mutant was included here as a tolerant control for reference. Mean root lengths of accessions grown on MV plates varied from 1.4 to 8.7 mm (Fig. S8b), indicating a wide variation in the MV response of *Arabidopsis*. Similarly, diverse responses were observed using the chlorophyll fluorescence assay; F_v_ F_m_^−1^ values varied from 0.109 in the sensitive Ag-0 ecotype to 0.694 in the tolerant Bil-7 (Fig. S8c). With few exceptions, the relative response to MV-induced ROS of these accessions under illuminated conditions was reproducible in all the assays above.

A set of accessions representing varied responses to organellar ROS were selected for further study (Fig. 5), including the relatively sensitive Kz-1, Col-0, Ga-0, and HR-10, the moderate Kondara, Ler, Zdr-1, Ws-2, Cvi-0, and Ll-0, and the relatively tolerant Mr-0, Lov-1, Bil-7 and the *rcd1* mutant. APX protein levels could not explain the observed natural variation in ROS sensitivity (Fig. S9). To test for differences, these accessions were assayed under both light and dark conditions (Fig. 5a,b). About half of these had similar sensitivity in both light and dark, while six genotypes changed in their relative sensitivity; Col-0, Cvi-0 and Kondara had increased tolerance in the dark while Mr-0, Ler and Ws-2 had greater sensitivity (Fig. 5a,b; in both panels the accessions are ordered according to tolerance under light). This demonstrates large natural variation in organellar ROS sensitivity also under dark conditions and suggests responses are conditioned by distinct loci in light and dark.

**Fig. 5.**
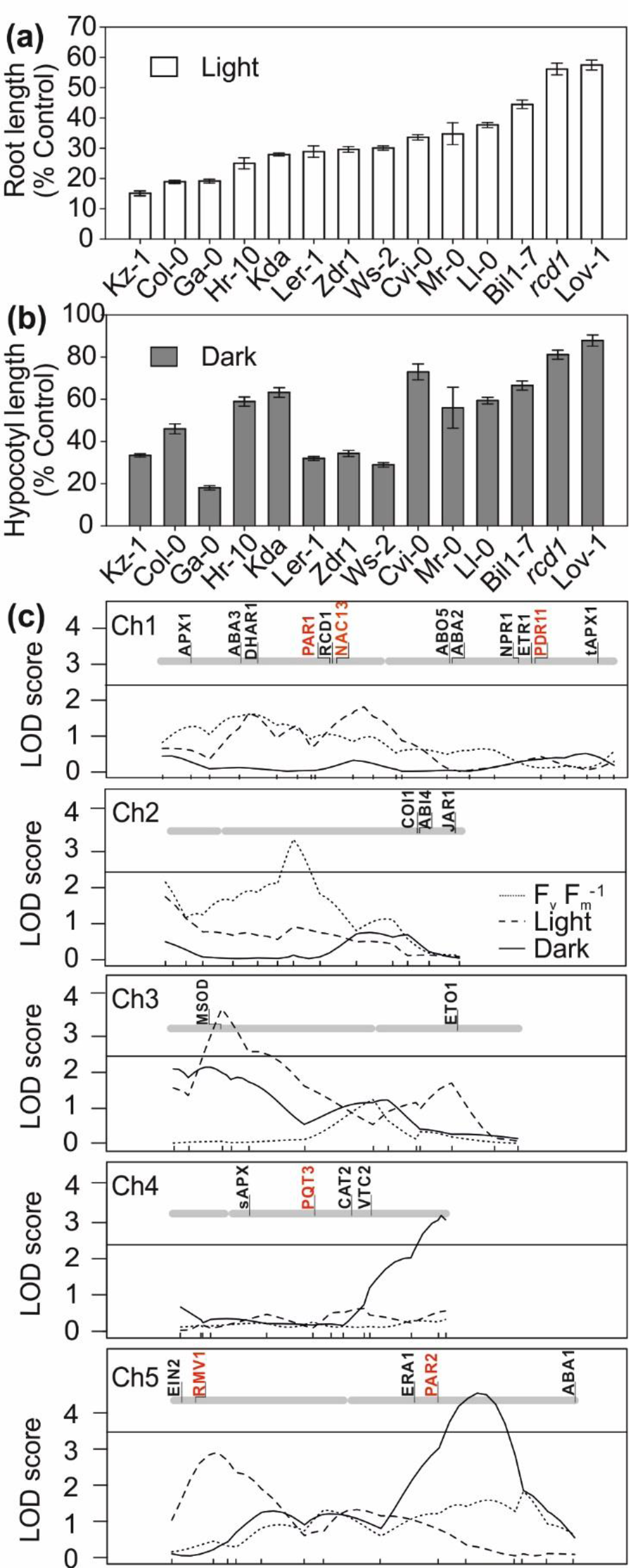
Natural variation in methyl viologen (MV)-induced ROS sensitivity in the light and dark. Roots lengths (a) or hypocotyl lengths (b) presented as percent of control of light or dark grown accessions in 0.1μM MV. Results are presented as means ±SD (N=33) of the ratio between MV treated to control root/hypocotyl lengths expressed as a percentage (% control). Experiment was repeated three times and results were pooled and analysed with posthoc analysis by computing contrasts from linear models and subjecting the P values to single-step error correction. Root lengths and hypocotyl were determined from eight or nine day old seedlings grown on 0.5x MS plates 1% sucrose and 0.1μM MV in the light (a) and dark (b) respectively. (c) Quantitative train loci (QTL) mapping in a Kondara×L*er* recombinant inbred line (RIL) population. Three separate MV traits were used: light root length (dashed lines); dark hypocotyl length (solid line) and chlorophyll fluorescence (quantum efficiency of photosystem II expressed as F_v_ F_m_^−1^; dotted lines). The genome-wide LOD threshold (horizontal line) for a QTL significance (P < 0.05) was calculated with 10 000 permutations and an average over the three traits is presented here (LOD = 2.4). QTL analysis was performed on the means of three biological repeats. Sample numbers were as follows; mapping in the light, N=12-20, mapping in the dark, N= 12-20, mapping with chlorophyll fluorescence, N=15-20. The positions of genes used in this study are indicated in black on top of the chromosomes; MV response genes previously identified by QTL mapping or forward genetics are indicated in red. For names, AGI codes, and references for the genes depicted in panel (c) see Table S3

An RIL population for the cross of L*er* and Kondara (El-Lithy *et al.*, 2006), whose relative MV-sensitivity changed between light and dark (Fig. 5a,b), was selected for in depth analysis in light and dark using QTL mapping with three different assays; chlorophyll fluorescence and root growth in the light and hypocotyl growth in the dark. In the chlorophyll fluorescence assay (F_v_ F_m_^−1^), one QTL was identified on the lower arm of chromosome two (Fig. 5c, dotted lines) and in the root growth assay in the light two additional QTLs were identified; one on the upper arm of chromosome three and one on the upper arm of chromosome five (Fig. 5c, dashed lines). Dark conditions revealed two additional distinct QTLs on the bottom of chromosome four and the lower arm of chromosome five (Fig. 5c, solid lines). All QTLs identified here were distinct from previously known MV-response QTLs (in red in Fig. 5c; gene list with AGI codes listed in Table S3; Fujita *et al.*, 2012). Taken together, these data suggest multiple mechanisms underpin the observed natural variation in organellar ROS tolerance, with distinct genetic loci regulating the responses in the light and dark.

### Stress hormones

To address the role of hormone signalling, a collection of 10 stress-hormone and ROS-signalling mutants were tested using growth assays under light and dark conditions. For a list of genotypes tested, mutant names, and AGI codes, see Table S4. The results (Fig. 6) are displayed in groups of functionally related mutants involved in salicylic acid (SA), jasmonic acid (JA), ethylene (ET), and ROS scavenging (Fig. 6a) and ABA (Fig. 6c). The organellar ROS sensitivity of the *vitamin c2*-1 (*vtc2*-1) mutant confirmed the role of ascorbate (ASC) in the light and to a lesser extent in the dark. Plants with diminished SA accumulation (*NahG*) displayed somewhat deficient tolerance both in the light and dark (Fig. 6a) implicating SA in organellar ROS signalling. In contrast, impaired ET-signalling led to minor tolerance. While ABA deficient mutants were mostly similar to wild type, mutants with enhanced ABA responses (*era1*-2) or ABA over-accumulation (*abo5*-2) were tolerant. The SA insensitive *npr1*-1, which hyper-accumulates SA, also exhibited tolerance. This suggests that hormone signalling- or metabolic-imbalances can modulate organellar ROS-induced sensitivity.

**Fig. 6.**
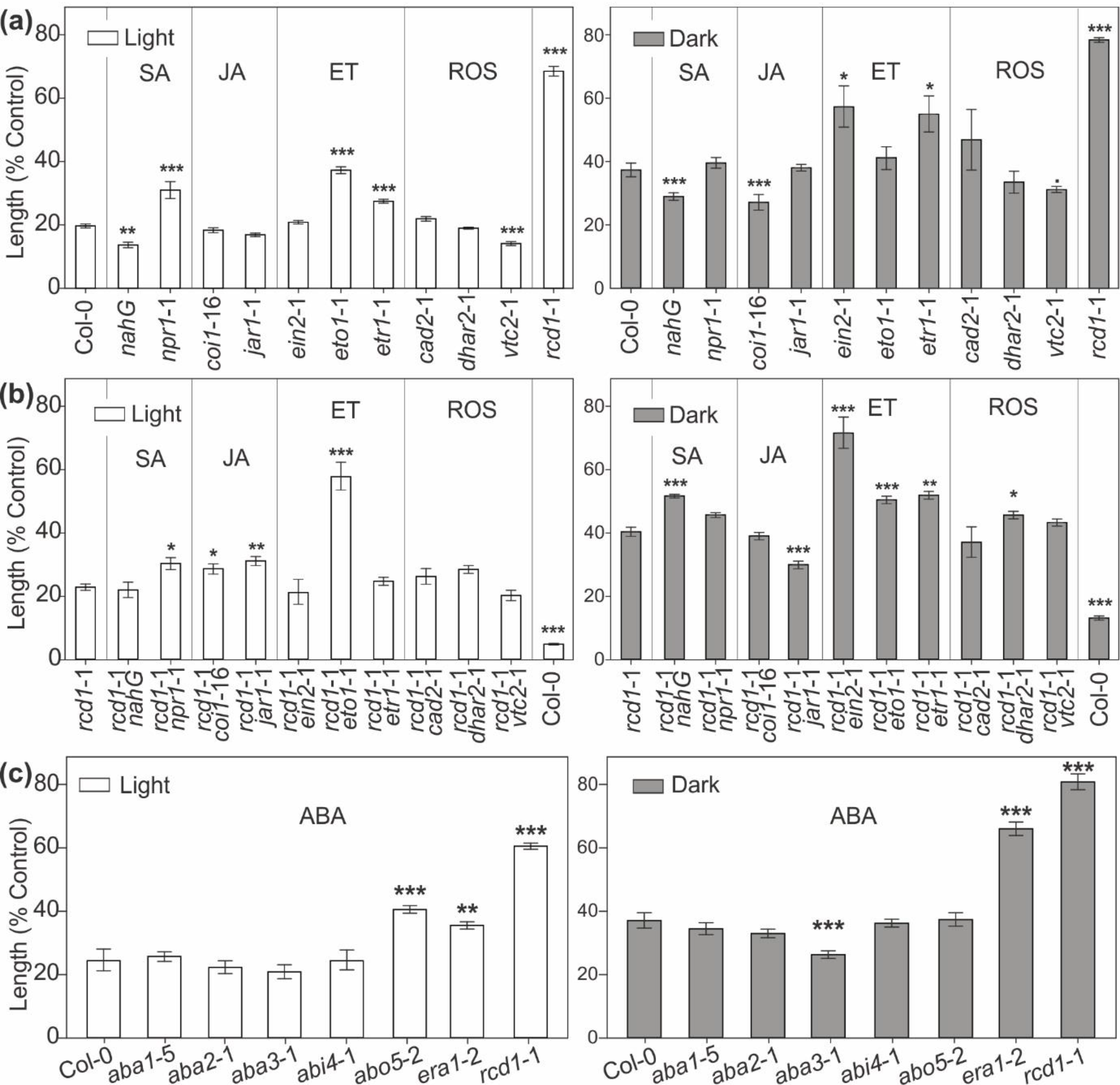
Methyl viologen (MV) response of hormone mutants in light and dark conditions. Reverse genetic experiments with single mutants related to salicylic acid (SA), jasmonic acid (JA), ethylene (ET) and reactive oxygen species (ROS) in (a). Results for *rcd1* double mutants are presented in (b) with root length assayed the light (left panel) and hypocotyl length in the dark (right panel). Results for abscisic acid (ABA) are presented in (c). Results are presented as means ±SD (N=32) of the ratio between MV treated to control (plants grown on identical plates without MV) lengths expressed as a percentage (% control). Experiment was repeated four times and results were pooled and analysed. Statistical significance was calculated from posthoc analysis by computing contrasts from linear models and subjecting the P values to single-step error correction. Measurements were from eight- or nine-day-old seedlings grown in the light or dark on 0.5x MS plates 1% sucrose and 0.2μM MV in panel (b) and 0.1μM MV in all other panels. P-value<0.01 ‘***’, P-value<0.01 ‘**’, P-value<0.05 ‘*’, P-value<0.1 ‘.’. List of genotypes tested, including full mutant names and AGI codes, is available in Table S4

To address hormone signalling in *rcd1* tolerance to MV-induced ROS, 10 *rcd1* double mutants (Overmyer *et al.*, 2000; Overmyer *et al.*, 2005; Blomster *et al.*, 2011; Brosché *et al.*, 2014) were assayed using higher (0.2 μM) MV to achieve similar relative growth inhibition in Col-0 and *rcd1* (Fig. 6b). The results were again organized into functionally related groups, as above. Increased tolerance was more common than sensitivity (Fig. 6b). The *rcd1 jar1*-1 mutant had opposite phenotypes in the light and dark, but the *jar1*-1 single mutant had a wild type phenotype. In the dark *jar1*-1 partially suppressed the *rcd1* tolerance phenotype, as *rcd1 jar1*-1 had reduced tolerance relative to the *rcd1*. In the light, *rcd1* double mutants with *eto1*-1, *coi1*-16, *jar1*-1 and *npr1*-1 exhibited further enhancement of tolerance. Similarly, many mutations further enhanced *rcd1* tolerance in the dark, including *rcd1* double mutants with *ein2-1, eto1-1, etr1-1*, and *NahG*.

The experiments above indicate a role for stress hormones, which we further tested using exogenous hormone treatment of photosynthetically active seedlings in the light using the chlorophyll fluorescence assay (Fig. 7). MV treatment resulted in visible symptoms at 24 hr (Fig. 7a) and decreased F_v_ F_m_^−1^ at six hr (Fig. 7b,c). Pre-treatment with ABA, SA or methyl jasmonate (JA), but not the ethylene precursor ACC, resulted in significant attenuation of MV damage. This could be seen both at the level of symptom development and F_v_ F_m_^−1^ (Fig. 7). These results further support the conclusions that the stress hormones ABA, SA and JA are regulators of plant MV-induced ROS tolerance. Hormone treatments were unable to induce further tolerance in the *rcd1* mutant (Fig. S10).

**Fig. 7.**
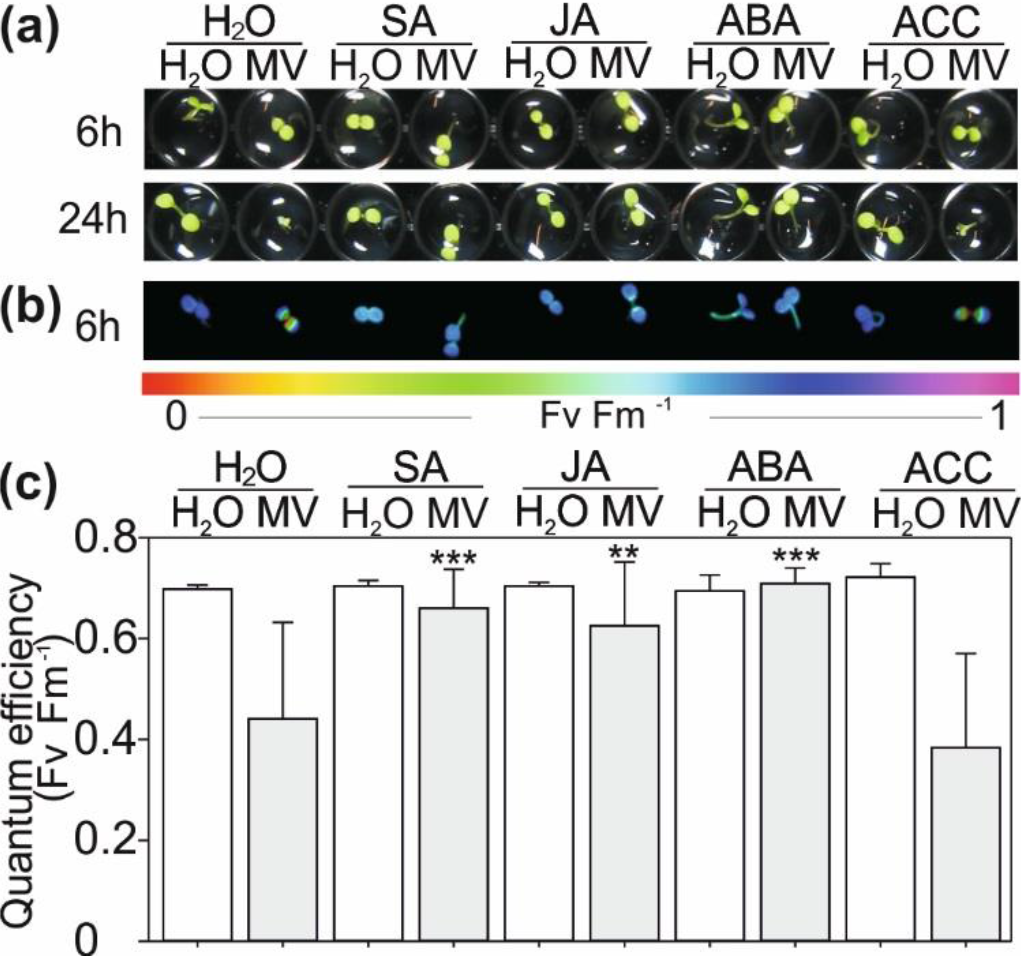
Protection from methyl viologen (MV) damage by phytohormones. (a) The phytohormones abscisic acid (ABA), salicylic acid (SA) and methyl jasmonic acid (JA), but not the ethylene precursor 1-aminocyclopropane-1-carboxylic acid (ACC), could protect plant from MV. Col-0 seedlings were pre-treated with hormones (0.2 mM) or water control (H_2_ O) for 12 hr before adding MV (0.25 mM) or water. Photos were taken 6 and 12 hr after MV treatment. Plants were one-week-old seedlings, grown one per well in a 96 well plate containing 180 μl of 0.5x MS media treated with 20 μl solutions containing MV. Experiment was repeated three times with similar results. One representative experiment is shown. (b) False colour image of quantum efficiency of photosystem II values (F_v_ F_m_^−1^) measured from chlorophyll fluorescence of seedlings treated as in (a). Chlorophyll fluorescence was measured with an Imaging PAM from seedlings 6 hr after MV treatment. Growth and treatment was as in (a). Experiment was repeated three times with similar results. One representative experiment is shown. (c) Quantification of quantum efficiency eight hr after MV treatment. Results are presented as means ±SD (N=24). Experiment repeated three times with similar results. One representative experiment is shown. Growth and treatment was as in (a)

## Discussion

### Mitochondria in MV-induced ROS signalling

Multiple genetic studies presented here support that MV could initiate ROS signals in the dark, when chloroplastic electron transfer is not active. MV responses in the light, when photosynthetic electron transport is active, were frequently different from those in the dark, suggesting that distinct signalling pathways control the light and dark response to MV-induced ROS. Hence, in addition to the classical light-dependent mechanism in the chloroplast, there is another ROS signalling pathway, as there is in non-photosynthetic organisms (Krall *et al.*, 1988; Minton *et al.*, 1990; Cochemé & Murphy, 2008), where MV induces ROS formation in the mitochondrial electron transfer chain. The site of MV action in the plant mitochondria should be addressed in future studies. In animals and yeast, MV acts to produce ROS at complex I, on the stromal side of the inner membrane (Cochemé and Murphy, 2008). It is conceivable that MV may act in the chloroplast in the dark. Some biochemical processes in the chloroplast also function in the dark, as seen in *Chlamydomonas* (Johnson & Alric, 2013; Cheung *et al.*, 2014). Further, the reduction of MV was observed in the dark in isolated chloroplasts (Law et al., 1983). However, the lack of MV-induced chloroplast stress in the dark (Fig 2) argues against this and supports the role of mitochondria in MV responses.

The potentiation of MV-induced ROS by exogenous sugar further implicates mitochondria in MV-triggered ROS signalling. Exogenous sugar enhanced MV-induced ROS responses in both light and dark suggesting that increased mitochondrial electron flow from activation of oxidative phosphorylation (Keunen *et al.*, 2013) potentiates MV-induced mitochondrial ROS. Sugars have tight connections to energy balance, redox balance, and ROS production due to their involvement in photosynthesis, oxidative phosphorylation and fatty acid beta-oxidation (Couée *et al.*, 2006; Keunen *et al.*, 2013). Furthermore, sugars are directly perceived and have dedicated signalling pathways to control and balance energy relations (Li & Sheen, 2016). These pathways are well integrated into several plant hormone signalling pathways, such as ethylene and ABA (Gazzarrini & McCourt, 2001). Thus, an alternative interpretation would be that sugars enhance ROS signalling by direct sugar-signalling pathways. We reasoned that if this were true, then the known sugar-hypersensitive hormone signalling mutants used here (*ein2*-1, *etr1*-1, *abo5*-2, and *era1*-2) should be MV sensitive, while sugar-insensitive mutants (*eto1*-1, *aba1*-1, *aba2*-1, *aba3*-1, and *abi4*-1) should be MV tolerant. This was not the case. Only the *eto1*-1 mutant behaved consistent with this model; all other sugar-signalling mutants exhibited WT responses or were opposite to the above predictions. This suggests that synergism of MV and exogenous sugar is independent of sugar signalling and rather supports the model where the exogenous sugar used in our experimental system activates oxidative phosphorylation and mitochondrial electron transport. Finally, lines lacking the mitochondrial MnSOD exhibited enhanced sensitivity in both light and dark, providing further evidence for mitochondria in MV-induced ROS signalling. The involvement of these mitochondrial processes in the MV-induced ROS response in the light, which was previously considered to involve only the chloroplast, suggests that chloroplast and mitochondrial ROS signalling pathways act in concert in response to MV. Furthermore, this suggests different partially overlapping MV-induced ROS signalling mechanisms in different situations; involving the mitochondria in the dark and the chloroplast and mitochondria in the light.

### ROS signalling and cytosolic ascorbate metabolism

Our results demonstrate that the role for ASC is dependent on its location. Knockouts of the chloroplast localized APXs (*tapx* and *sapx*), residing near the site of chloroplast ROS production (Asada, 1999) under light conditions had normal MV-induced ROS phenotypes (Fig. S7c) and photosynthesis rates unchanged from wild type under moderate light stress (1000 μmol m^−2^ s^−1^ illumination; Davletova *et al.*, 2005). This seemingly counterintuitive result may be explained by the multiple effects MV has on chloroplasts. MV competes with ferredoxin for electrons at PSI, resulting in ROS production, but also diverting electrons from ferredoxin and its downstream electron acceptors. Accordingly, MV-treatment results in a decrease in the NADPH pool (Benina *et al.*, 2015), the rapid oxidation of chloroplast ASC and GSH, and the disappearance of dehydroascorbate (Law *et al.*, 1983). Thus, MV treatment results in attenuation of chloroplast protective pathways such as the water-water cycle, cyclic electron transport, and the ASC-glutathione (GSH) cycle (Law *et al.*, 1983; Hanke & Mulo, 2013). Together these results suggest the existence of chloroplast protective pathways that either divert electron flow to reduce ROS production or derive reducing power for ROS detoxification from sources other than PSI. The ASC deficient *vtc2* (Fig. 6) mutant and cytosolic *capx* mutants were MV sensitive in the light and dark (*capx1-2*, Fig. S7; *capx1-1*, Davletova *et al.*, 2005), suggesting a role for cytosolic ASC. Previously, MV-treatment was shown to result in the accumulation of cytosolic H_2_O_2_ (Schwarzländer *et al.*, 2009). Also, a requirement for cytosolic APX to maintain normal photosynthesis rates under illumination of 1000 μmol m^−2^ s^−1^ was demonstrated (Davletova *et al.*, 2005). This involvement of a cytosolic ROS scavenger for chloroplast protection suggests complex inter-compartmental signalling. Indeed, the *capx1-1* mutant was previously shown to have altered transcriptional profiles for many signalling genes and redox modifications of several key signalling proteins (Davletova *et al.*, 2005). This suggests that cAPX modulates ROS in the regulation of an inter-compartmental signalling pathway involving both photosynthetic and non-photosynthetic mechanisms. Taken together, our results support a model where ROS signalling pathways from both inside and outside the chloroplast determine the plant response to MV (Fig. 8).

**Fig. 8.**
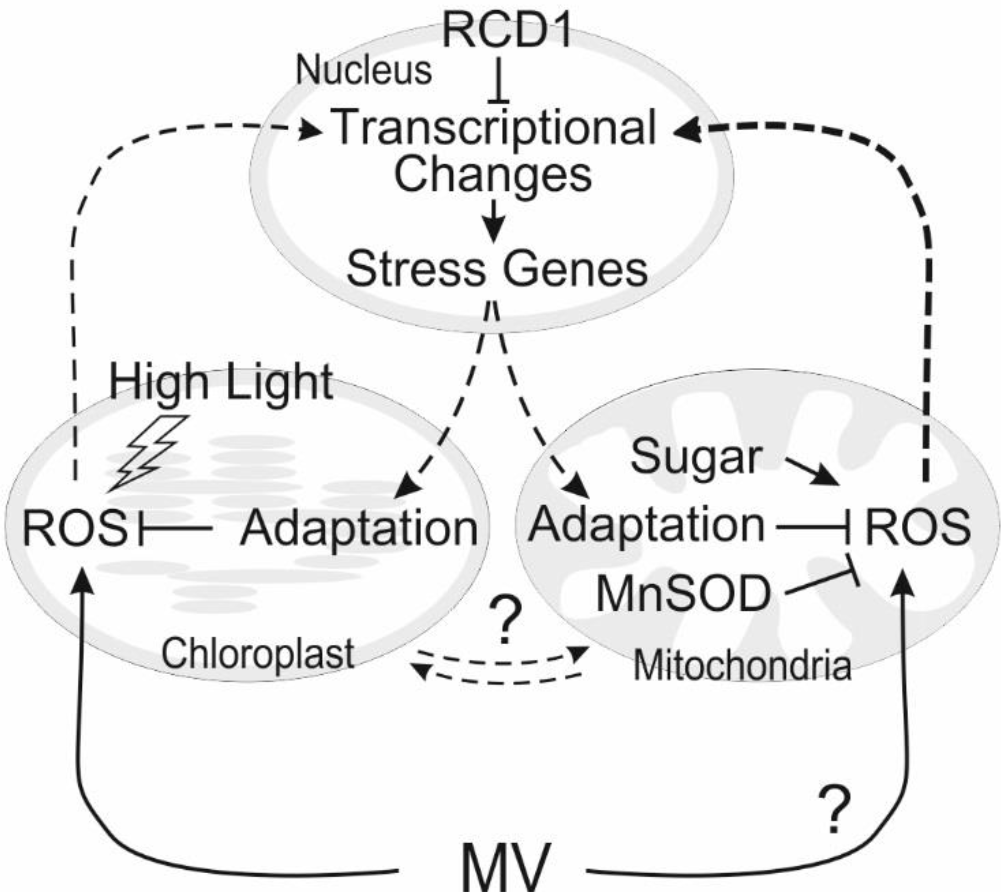
Model of methyl viologen (MV)-induced ROS signalling. Diagram depicting a proposed signalling network where MV-induced ROS formation in either the mitochondria or chloroplast results in ROS signals that trigger retrograde signalling back to the nucleus, which in turn activates stress responsive transcriptional programs responsible for adaptive responses in both organelles. Lines with a question mark indicate two processes suggested but not proven by the data in this work; MV-induced mitochondrial ROS formation and the nature of the functional link between the mitochondria and chloroplast. RADICAL-INDUCED CELL DEATH1 (RCD1), Manganese (mitochondrial) SUPEROXIDE DISMUTASE (MnSOD)

### The role of stress hormones

Our results implicated the plant stress hormones in the organellar ROS response (Fig. 6). Results with SA-deficient (*NahG*) and SA-hyper-accumulating (*npr1*) plants suggest SA modulates intercellular ROS signalling in an NPR1-independent manner. SA is a known inhibitor of mitochondrial electron transport and inducer of mitochondrial dysfunction stimulon (MDS) marker genes (Norman *et al.*, 2004; Van Aken *et al.*, 2009), consistent with the role for mitochondria proposed here. JA signalling has been implicated in chloroplast retrograde signalling (Tikkanen *et al.*, 2014), but may also act indirectly via its mutually antagonistic interaction with SA. Also, ET modulates the xanthophyll cycle to increase ROS production and photosensitivity by the suppression of non-photochemical quenching (Chen & Gallie, 2015). Accordingly, exogenous JA, ABA, and SA treatments induce tolerance to MV-induced ROS in Col-0 (Fig. 7). These hormones do not confer any additional tolerance to *rcd1* mutant plants, suggesting that hormone-signalling and RCD1-dependent ROS signalling converge into a common downstream pathway that modulates protective responses.

In root-growth assays, ET signalling single and double mutants enhanced ROS tolerance, demonstrating additive effects in long-term developmental responses over the course of days. However, treatment of Col-0 plants with exogenous the ET precursor, ACC, had no additional effect, as measured in short term experiments lasting hours using the chlorophyll fluorescence assay. This is likely due to differences in the assays used. Several of our experiments demonstrate variability in the MV-response dependent on growth conditions and the assay used (Figs. 2, 3, and 6; Fig. S2; Fig. S3). This was especially apparent in the QTL mapping (Fig. 5; Fig. S8), where different QTLs were identified depending on the assay used, illustrating that different assays can detect distinct genetic pathways governing the MV-induced ROS response. Thus, caution must be exercised in comparing results between experiments using different assays.

### RCD1 and retrograde signalling

RCD1 acts on multiple ROS signalling pathways in distinct subcellular compartments, including stress protection pathways (Shapiguzov *et al.*, 2018). RCD1 is a plant-specific protein that interacts with a specific set of transcription factors regulating multiple stress- and developmental-pathways (Ahlfors *et al.*, 2004; Jaspers *et al.*, 2009; Vainonen *et al.*, 2012). Analysis of RCD1-regulated genes revealed many misregulated MDS genes (Jaspers *et al.*, 2009; Brosché *et al.*, 2014), which are markers of mitochondrial retrograde regulation (MRR) signalling, suggesting that RCD1 is involved in the transmission of ROS signals from the mitochondria to the nucleus. High-level overexpression of mitochondrial dysfunction stimulon (MDS) genes in *rcd1* indicates that RCD1 is also involved in retrograde signalling that results in mitochondrial stress adaptation. Our results show RCD1-dependent alterations in both chloroplasts and mitochondria, suggesting coordinated responses between the two organelles (Fig. 8), accordingly the *rcd1* mutant was highly tolerant of MV-induced ROS in both the light and dark. However the question remains, from which organelle does the primary effect on MV-induced ROS responses originate? Two lines of evidence support that RCD1 is a regulator of primarily mitochondrial processes. First, there is a large difference in magnitude between the high-light and MV phenotypes in the *rcd1* mutant; the weaker phenotype is high light stress, which is a purely chloroplastic stress. Further, genes deregulated in the *rcd1* mutant showed significant enrichment for genes encoding proteins residing in the mitochondria, but not in the chloroplast. Together these findings support a model where RCD1 acts primarily through the mitochondria to modulate MV-induced ROS signalling.

The MRR regulators, NO APICAL MERISTEM/ARABIDOPSIS TRANSCRIPTION ACTIVATION FACTOR/CUP-SHAPED COTYLEDON13 (ANAC013) and ANAC017 transcription factors (De Clercq *et al.*, 2013; Ng *et al.*, 2013), are among the transcription factors that interact with RCD1 (Jaspers *et al.*, 2009; Shapiguzov *et al.*, 2018). ANAC013-overexpression enhanced tolerance to MV-induced ROS (De Clercq *et al.*, 2013) when assayed for visual symptoms (leaf bleaching and chlorosis), leaf fresh weight and root growth in the light using 0.1 μM MV, the same as in the current study. This suggests either that ANAC013 directly regulates genes important for proper chloroplast function, or an indirect interaction between the mitochondria and chloroplast. Similarly, this concept has been seen before, the ABI4 transcription factor is involved in both chloroplast and mitochondrial retrograde signalling (León *et al.*, 2013; Giraud *et al.*, 2009). MDS and MRR genes are positively regulated by ANAC013 and their expression is negatively regulated by RCD1; supporting RCD1 as a regulator of MRR via its negative regulation of ANAC013 function. In a related study, ROS signals from the mitochondria and chloroplast were shown to converge on the redox regulation of RCD1 (Shapiguzov *et al.*, 2018) to alter the expression of MDR genes including alternative oxidases (*AOXs*). Enhanced accumulation of these MDR genes altered chloroplastic electron flow, decreasing chloroplastic ROS and associated damage (Shapiguzov *et al.*, 2018).

Taken together our results support the role of mitochondrial processes in the MV response. We propose that interactions between the chloroplast and mitochondria, regulated by RCD1 and stress hormones, are involved in determining plant response to redox imbalance during MV treatment.

## Acknowledgments

We thank Tuomas Puukko and Leena Grönholm for excellent technical assistance; Eva-Mari Aro for PSI and PSII subunit antibodies; Patricia Conklin (*vtc2*), Lee Sweetlove (MnSOD RNAi line) and Zhizhong Gong (*abo5*) for seeds; and Mohamed E. El-Lithy and Martin Koornneef for the Kondara x L*er* RIL genotyping data. This work was supported by the University of Helsinki (three-year research allocations to M.B. and K.O.) and Academy of Finland Fellowships (Decisions no.135751, 140981 and 273132 to M.B., no. 251397, 256073, and 283254 to K.O. and no. 263772, 218157, 259888 and 130595 to SK). KO, JS, JK, MB, and SK were supported by the Academy of Finland Center of Excellence in Molecular Biology of Primary Producers 2014-2019 (Decisions no. 307335 and 271832). FC was a member of the University of Helsinki Doctoral Program in Plant Sciences (DPPS).

## Supporting Information

**Fig. S1** Transcript accumulation of known methyl-viologen (MV) transporter genes.

**Fig. S2** Organellar ROS tolerance of four *radical-induced cell death1* (*rcd1*) alleles.

**Fig. S3** Diaminobenzidine (DAB) staining of methyl viologen (MV)-induced H_2_ O_2_ accumulation in the dark.

**Fig. S4** Glucose enhanced the methyl-viologen (MV)-induced ROS response in light and dark.

**Fig. S5** Optimization of assay conditions for genetic studies.

**Fig. S6** Chloroplast adaptation to fluctuating light stress in Col-0 and *radical-induced cell death1* (*rcd1*) mutant plants.

**Fig. S7** Ascorbate peroxidase (APX) mutants.

**Fig. S8** Natural variation in the Arabidopsis methyl viologen (MV)-response.

**Fig. S9** Ascorbate peroxidase protein levels in Arabidopsis natural accessions under control and methyl viologen (MV) treated conditions.

**Fig. S10** Effect of exogenous stress hormone treatment on methyl viologen (MV) tolerance of the *radical-induced cell death1* (*rcd1*) mutant.

**Table S1** Primers used in this study.

**Table S2** Information about genes used in Figure 3.

**Table S3** Information about genes used in Figure 5c.

**Table S4** Mutants used in this study.

## References

Abarca D, Roldán M, Martín M, and Sabater, B. 2001. Arabidopsis thaliana ecotype Cvi shows an increased tolerance to photo‐oxidative stress and contains a new chloroplastic copper/zinc superoxide dismutase isoenzyme. Journal of Experimental Botany 52: 1417–1425.

Ahlfors R, Lång S, Overmyer K, Jaspers P, Brosché M, Tauriainen A, Kollist H, Tuominen H, Belles-Boix E, Piippo M et al. 2004. Arabidopsis RADICAL-INDUCED CELL DEATH1 belongs to the WWE protein-protein interaction domain protein family and modulates abscisic acid, ethylene, and methyl jasmonate responses. Plant Cell 16: 1925–1937.

Aono, M, Saji H, Sakamoto A, Tanaka K, Kondo N, Tanaka K. 1995. Paraquat tolerance of transgenic Nicotiana tabacum with enhanced activities of glutathione reductase and superoxide dismutase. Plant and Cell Physiology 36: 1687–1691.

Arnon DI. 1949. Copper enzymes in isolated chloroplasts. Polyphenoloxidase in Beta vulgaris. Plant Physiology 24: 1–15.

Asada K. 1999. The water-water cycle in chloroplasts: scavenging of active oxygens and dissipation of excess photons. Annual Review of Plant Physiology and Plant Molecular Biology 50: 601–639.

Babbs CF, Pham JA, Coolbaugh, RC. 1989. Lethal hydroxyl radical production in paraquat-treated plants. Plant Physiology 90: 1267–1270.

Barbagallo RP, Oxborough K, Pallett KE, Baker NR. 2003. Rapid, noninvasive screening for perturbations of metabolism and plant growth using chlorophyll fluorescence imaging. Plant Physiology 132: 485–493.

Belles-Boix E, Babiychuk E, Van Montagu M, Inzé D, Kushnir S. 2000. CEO1, a new protein from Arabidopsis thaliana, protects yeast against oxidative damage. FEBS Letters 482: 19–24.

Benina M, Mendes Ribeiro D, Gechev TS, Mueller-Roeber B, Schippers JHM. 2015. A cell type specific view on the translation of mRNAs from ROS-responsive genes upon paraquat treatment of Arabidopsis thaliana leaves. Plant Cell and Environment 38: 349–363.

Bowler C, Slooten L, Vandenbranden S, De Rycke R, Botterman J, Sybesma C, Van Montagu M, Inzé D. 1991. Manganese superoxide dismutase can reduce cellular damage mediated by oxygen radicals in transgenic plants. EMBO Journal 10: 1723–1732.

Bretz F, Hothorn T, Westfall P. 2010. Multiple comparisons using R. Boca Raton, FL: CRC Press.

Blomster T, Salojärvi J, Sipari N, Brosché M, Ahlfors R, Keinänen M, Overmyer K, Kangasjärvi J. 2011. Apoplastic reactive oxygen species transiently decrease auxin signaling and cause stress-induced morphogenic response in Arabidopsis. Plant Physiology 157: 1866–1883

Broman KW, Wu H, Sen S, Churchill GA. 2003. R/qtl: QTL mapping in experimental crosses. Bioinformatics 19: 889–890.

Brosché M, Blomster T, Salojärvi J, Cui F, Sipari N, Leppälä J, Lamminmäki A, Tomai G, Narayanasamy S, Reddy RA et al. 2014. Transcriptomics and functional genomics of ROS-induced cell death regulation by RADICAL-INDUCED CELL DEATH1. PLoS Genetics. doi: 10.1371/journal.pgen.1004112.

Chen R, Sun S, Wang C, Li Y, Liang Y, An F, Li C, Dong H, Yang X, Zhang J et al. 2009. The Arabidopsis PARAQUAT RESISTANT2 gene encodes an S-nitrosoglutathione reductase that is a key regulator of cell death. Cell Research 19: 1377–1387.

Chen Z, Gallie DR. 2015. Ethylene regulates energy-dependent non-photochemical quenching in Arabidopsis through repression of the xanthophyll cycle. PLoS ONE. doi.org/10.1371/journal.pone.0144209.

Cheung CYM, Poolman MG, Fell DA, Ratcliffe RG, Sweetlove LJ. 2014. A diel flux balance model captures interactions between light and dark metabolism during day-night cycles in C3 and crassulacean acid metabolism leaves. Plant Physiology 165: 917–929.

Cochemé HM, Murphy MP. 2008. Complex I is the major site of mitochondrial superoxide production by paraquat. Journal of Biological Chemistry 283: 1786–1798.

Couée I, Sulmon C, Gouesbet G, El Amrani A. 2006. Involvement of soluble sugars in reactive oxygen species balance and responses to oxidative stress in plants. Journal of Experimental Botany 57: 449–459.

Davletova S, Rizhsky L, Liang H, Shengqiang Z, Oliver DJ, Coutu J, Shulaev V, Schlauch K, Mittler R. 2005. Cytosolic ascorbate peroxidase 1 is a central component of the reactive oxygen gene network of Arabidopsis. Plant Cell 17: 268–281.

De Clercq I, Vermeirssen V, Van Aken O, Vandepoele K, Murcha MW, Law SR, Inzé A, Ng S, Ivanova A, Rombaut D et al. 2013. The membrane-bound NAC transcription factor ANAC013 functions in mitochondrial retrograde regulation of the oxidative stress response in Arabidopsis. Plant Cell 25: 3472–3490.

Dodge AD. 1989. Herbicides interacting with photosystem I. In: Dodge A.D., ed. Herbicides and plant metabolism, Society for Experimental Biology seminar series. Cambridge, UK, Cambridge University Press, 1–277.

El-Lithy ME, Bentsink L, Hanhart CJ, Ruys GJ, Rovito D, Broekhof JLM, van der Poel HJA, van Eijk MJT, Vreugdenhil D, Koornneef M. 2006. New Arabidopsis recombinant inbred line populations genotyped using SNPWave and their use for mapping flowering-time quantitative trait loci. Genetics 172: 1867–1876.

Fernie AR, Carrari F, Sweetlove LJ. 2004. Respiratory metabolism: glycolysis, the TCA cycle and mitochondrial electron transport. Current Opinion Plant Biology 7: 254–261.

Foyer CH, Noctor G. 2011. Ascorbate and glutathione: the heart of the redox hub. Plant Physiology 155: 2–18.

Foyer CH, Noctor G. 2016. Stress-triggered redox signalling: what’s in pROSpect? Plant Cell and Environment 39: 951–964.

Fuerst EP, Norman MA. 1991. Interactions of herbicides with photosynthetic electron transport. Weed Science 39: 458–464.

Fujibe T, Saji H, Arakawa K, Yabe N, Takeuchi Y, Yamamoto KT. 2004. A methyl viologen-resistant mutant of Arabidopsis, which is allelic to ozone-sensitive rcd1, is tolerant to supplemental ultraviolet-B irradiation. Plant Physiology 134: 275–285.

Fujita M, Fujita Y, Iuchi S, Yamada K, Kobayashi Y, Urano K, Kobayashi M, Yamaguchi-Shinozaki K, Shinozaki K. 2012. Natural variation in a polyamine transporter determines paraquat tolerance in Arabidopsis. Proceedings of the National Academy of Sciences of the United States of America 109: 6343–6347.

Fujita M, Shinozaki K. 2014. Identification of polyamine transporters in plants: paraquat transport provides crucial clues. Plant and Cell Physiology 55: 855–861.

Gazzarrini S, McCourt P. 2001. Genetic interactions between ABA, ethylene and sugar signaling pathways. Current Opinion in Plant Biology 4: 387–391.

Giraud E, Van Aken O, Ho LHM, Whelan J. 2009. The transcription factor ABI4 is a regulator of mitochondrial retrograde expression of ALTERNATIVE OXIDASE1a. Plant Physiology 150: 1286–1296.

Hahn A, Kilian J, Mohrholz A, Ladwig F, Peschke F, Dautel R, Harter K, Berendzen KW, Wanke D. 2013. Plant core environmental stress response genes are systemically coordinated during abiotic stresses. International Journal of Molecular Sciences 14: 7617–7641.

Hanke G, Mulo P. 2013. Plant type ferredoxins and ferredoxin-dependent metabolism. Plant, Cell & Environment 36: 1071–1084.

Hassan HM. 1984. Exacerbation of superoxide radical formation by Paraquat. Methods in Enzymology 105: 523–532.

Hawkes TR. 2014. Mechanisms of resistance to paraquat in plants. Pest Management Science 70: 1316–1323.

Hruz T, Laule O, Szabo G, Wessendorp F, Bleuler S, Oertle, L, Widmayer P, Gruissem W, Zimmermann P. 2008. Genevestigator v3: a reference expression database for the meta-analysis of transcriptomes. Advances in Bioinformatics. doi: 10.1155/2008/420747.

Jaspers P, Blomster T, Brosché M, Salojärvi J, Ahlfors R, Vainonen JP, Reddy RA, Immink R, Angenent G, Turck F et al. 2009. Unequally redundant RCD1 and SRO1 mediate stress and developmental responses and interact with transcription factors. Plant Journal 60: 268–279.

Jaspers P, Overmyer K, Wrzaczek M, Vainonen JP, Blomster T, Salojärvi J, Reddy RA, Kangasjärvi J. 2010. The RST and PARP-like domain containing SRO protein family: analysis of protein structure, function and conservation in land plants. BMC Genomics. doi: 10.1186/1471-2164-11-170.

Johnson X, Alric J. 2013. Central carbon metabolism and electron transport in Chlamydomonas reinhardtii: metabolic constraints for carbon partitioning between oil and starch. Eukaryotic Cell 12: 776–793.

Schwarzländer M, Fricker MD, Sweetlove LJ. 2009. Monitoring the in vivo redox state of plant mitochondria: Effect of respiratory inhibitors, abiotic stress and assessment of recovery from oxidative challenge. Biochimica et Biophysica Acta 1787: 468–475.

Keunen E, Peshev D, Vangronsveld J, Van Den Ende W, Cuypers A. 2013. Plant sugars are crucial players in the oxidative challenge during abiotic stress: extending the traditional concept. Plant Cell and Environment 36: 1242–1255.

Krall J, Bagley AC, Mullenbach GT, Hallewell RA, Lynch RE. 1988. Superoxide mediates the toxicity of paraquat for cultured mammalian cells. Journal of Biological Chemistry 263: 1910–1914.

Kurepa J, Smalle J, Van Montagu M, Inzé D. 1998a. Oxidative stress tolerance and longevity in Arabidopsis: the late-flowering mutant gigantea is tolerant to paraquat. Plant Journal 14: 759–764.

Kurepa J, Smalle J, Van Montagu M, Inzé D. 1998b. Effects of sucrose supply on growth and paraquat tolerance of the late-flowering gi-3 mutant. Plant Growth Regululation 26: 91–96.

Lasat MM, DiTomaso JM, Hart JJ, Kochian LV. 1997. Evidence for vacuolar sequestration of paraquat in roots of a paraquat-resistant Hordeum glaucum biotype. Physiologia Plantarum 99: 255–262.

Law MY, Charles SA, Halliwell B. 1983. Glutathione and ascorbic acid in spinach (Spinacia oleracea) chloroplasts. Biochemical Journal 210: 899–903.

León P, Gregorio J, Cordoba E. 2013. ABI4 and its role in chloroplast retrograde communication. Frontiers in Plant Science. doi: 10.3389/fpls.2012.00304.

Li J, Mu J, Bai J, Fu F, Zou T, An F, Zhang J, Jing H, Wang Q, Li Z et al. 2013. Paraquat resistant1, a golgi-localized putative transporter protein, is involved in intracellular transport of paraquat. Plant Physiology 162: 470–483.

Li L, Sheen J. 2016. Dynamic and diverse sugar signaling. Current Opinion in Plant Biology 33: 116–125.

Luo C, Cai X-T, Du J, Zhao T-L, Wang P-F, Zhao P-X, Liu R, Xie Q, Cao X-F, Xiang C-B 2016. PARAQUAT TOLERANCE3 is an E3 ligase that switches off activated oxidative response by targeting histone-modifying PROTEIN METHYLTRANSFERASE4b. PLoS Genetics. doi.org/10.1371/journal.pgen.1006332.

Miller G, Suzuki N, Rizhsky L, Hegie A, Koussevitzky S, Mittler R. 2007. Double mutants deficient in cytosolic and thylakoid ascorbate peroxidase reveal a complex mode of interaction between reactive oxygen species, plant development, and response to abiotic stresses. Plant Physiology 144: 1777–1785.

Minton KW, Tabor H, Tabor CW. 1990. Paraquat toxicity is increased in Escherichia coli defective in the synthesis of polyamines. Proceedings of the National Academy of Sciences of the United States of America 87: 2851–2855.

Morgan MJ, Lehmann M, Schwarzländer M, Baxter CJ, Sienkiewicz-Porzucek A, Williams TCR, Schauer N, Fernie AR, Fricker MD, Ratcliffe RG et al. 2008. Decrease in manganese superoxide dismutase leads to reduced root growth and affects tricarboxylic acid cycle flux and mitochondrial redox homeostasis. Plant Physiology 147: 101–114.

Murgia I, Tarantino D, Vannini C, Bracale M, Carravieri S, Soave C. 2004. Arabidopsis thaliana plants overexpressing thylakoidal ascorbate peroxidase show increased resistance to Paraquat-induced photooxidative stress and to nitric oxide-induced cell death. Plant Journal 38: 940–953.

Ng S, Ivanov A, Duncan O, Law SR, Van Aken O, De Clercq I, Wang Y, Carrie C, Xu L, Kmiec B et al. 2013. A membrane-bound NAC transcription factor, ANAC017, mediates mitochondrial retrograde signaling in Arabidopsis. Plant Cell 25: 3450–3471.

Noguchi K, Yoshida K. 2008. Interaction between photosynthesis and respiration in illuminated leaves. Mitochondrion 8: 87–99.

Nordborg M, Hu TT, Ishino Y, Jhaveri J, Toomajian C, Zheng H, Bakker E, Calabrese P, Gladstone J, Goyal R et al. 2005. The pattern of polymorphism in Arabidopsis thaliana. PLoS Biology. doi:10.1371/journal.pbio.0030196

Norman C, Howell KA, Millar AH, Whelan JM, Day DA. 2004. Salicylic acid is an uncoupler and inhibitor of mitochondrial electron transport. Plant Physiology 134: 492–501.

Overmyer K, Tuominen H, Kettunen R, Betz C, Langebartels C, Sandermann H, Kangasjärvi J. 2000. Ozone-sensitive Arabidopsis rcd1 mutant reveals opposite roles for ethylene and jasmonate signaling pathways in regulating superoxide-dependent cell death. Plant Cell 12: 1849–1862.

Overmyer K, Brosché M, Pellinen R, Kuittinen T, Tuominen H, Ahlfors R, Keinänen M, Saarma M, Scheel D, Kangasjärvi J. 2005. Ozone-induced programmed cell death in the Arabidopsis radical-induced cell death1 mutant. Plant Physiology 137: 1092–1104.

R Development Core Team. 2014. R: A language and environment for statistical computing. [WWW document] URL http://www.R-project.org/. [accessed 1 May 2014]

Shapiguzov A, Vainonen JP, Hunter K, Tossavainen H, Tiwari A, Järvi S, Hellman M, Wybouw B, Aarabi F, Alseekh S et al. 2018. RCD1 coordinates chloroplastic and mitochondrial electron transfer through interaction with ANAC transcription factors. BioRXiv. doi.org/10.1101/327411

Tikkanen M, Grieco M, Kangasjärvi S, Aro E-M. 2010. Thylakoid protein phosphorylation in higher plant chloroplasts optimizes electron transfer under fluctuating light. Plant Physiology 152: 723–735.

Tikkanen M, Gollan PJ, Mekala NR, Isojärvi J, Aro E-M. 2014. Light-harvesting mutants show differential gene expression upon shift to high light as a consequence of photosynthetic redox and reactive oxygen species metabolism. Philosophical Transactions of the Royal Society London, B, Biological Sciences. doi: 10.1098/rstb.2013.0229

Vainonen JP, Jaspers P, Wrzaczek M, Lamminmäki A, Reddy RA, Vaahtera L, Brosché M, Kangasjärvi J. 2012. RCD1-DREB2A interaction in leaf senescence and stress responses in Arabidopsis thaliana. Biochemical Journal 442: 573–581.

Van Aken O, Giraud E, Clifton R, Whelan J. 2009. Alternative oxidase: a target and regulator of stress responses. Physiologia Plantarum 137: 354–361.

Van Camp W, Capiau K, Van Montagu M, Inze D, Slooten L. 1996. Enhancement of oxidative stress tolerance in transgenic tobacco plants overproducing Fe-superoxide dismutase in chloroplasts. Plant Physiology 112: 1703–1714.

Vanlerberghe G C, Martyn G D, Dahal K. 2016. Alternative oxidase: a respiratory electron transport chain pathway essential for maintaining photosynthetic performance during drought stress. Physiologia Plantarum 157: 322–337.

Váradi G, Darkó É, Lehoczki E. 2000. Changes in the xanthophyll cycle and fluorescence quenching indicate light-dependent early events in the action of paraquat and the mechanism of resistance to paraquat in Erigeron canadensis (L.) Cronq. Plant Physiology 123: 1459–1470.

Vaughn KC, Vaughan MA, Camilleri P. 1989. Lack of cross-resistance of paraquat-resistant hairy fleabane (Conyza bonariensis) to other toxic oxygen generators indicates enzymatic protection is not the resistance mechanism. Weed Science 37: 5–11.

Xi J, Xu P, Xiang C-B. 2012. Loss of AtPDR11, a plasma membrane-localized ABC transporter, confers paraquat tolerance in Arabidopsis thaliana. Plant Journal. 69: 782–791.

Yu Q, Cairns A, Powles SB. 2004. Paraquat resistance in a population of Lolium rigidum. Functional Plant Biology 31: 247–254.

Waszczak C, Carmody M, Kangasjärvi J. 2018. Reactive oxygen species in plant signaling. Annual Review of Plant Biology. doi.org/10.1146/annurev-arplant-042817-040322

